# Bayesian inference of ancestral dates on bacterial phylogenetic trees

**DOI:** 10.1101/347385

**Authors:** Xavier Didelot, Nicholas J Croucher, Stephen D Bentley, Simon R Harris, Daniel J Wilson

## Abstract

The sequencing and comparative analysis of a collection of bacterial genomes from a single species or lineage of interest can lead to key insights into its evolution, ecology or epidemiology. The tool of choice for such a study is often to build a phylogenetic tree, and more specifically when possible a dated phylogeny, in which the dates of all common ancestors are estimated. Here we propose a new Bayesian methodology to construct dated phylogenies which is specifically designed for bacterial genomics. Unlike previous Bayesian methods aimed at building dated phylogenies, we consider that the phylogenetic relationships between the genomes have been previously evaluated using a standard phylogenetic method, which makes our methodology much faster and scalable. This two-steps approach also allows us to directly exploit existing phylogenetic methods that detect bacterial recombination, and therefore to account for the effect of recombination in the construction of a dated phylogeny. We analysed many simulated datasets in order to benchmark the performance of our approach in a wide range of situations. Furthermore, we present applications to three different real datasets from recent bacterial genomic studies. Our methodology is implemented in a R package called BactDating which is freely available for download at https://github.com/xavierdidelot/BactDating.

## INTRODUCTION

A population evolving sufficiently quickly over a sufficiently long sampling time frame is said to be “measurably evolving”, which means that it is possible to estimate the rates over time at which evolution operates and the dates at which ancestors existed (1). This concept has recently become applicable to bacterial species, following the advent of whole-genome sequencing data, in which the relatively low per site evolutionary rates in bacteria are compensated by long genomes, typically comprising millions of sites (2). Consequently, analytical methods that were previously the hallmark of viral genetics are growing in popularity in bacterial genetics, especially the estimation of dated genealogies through the application of the software BEAST (3, 4, 5). In a dated phylogeny (also sometimes known as a time-stamped phylogeny or time-calibrated phylogeny), the branch lengths are measured in unit of time (for example days or years), the leaves are shown at known dates of isolation, and the internal nodes are represented at the dates when common ancestors are estimated to have existed. Such estimation of ancestral dates can often provide direct biological insights, for example to date the emergence of an epidemiologically important lineage, but can also be used as a starting point for further analysis, for example to infer past population size dynamics (6), to reconstruct transmission events between hosts (7), to estimate the parameters of an epidemiological model (8), to investigate geographical range expansion (9) or to study ecological adaptation to host species (10).

The BEAST framework is popular because it includes many models and extensions, and is based on the Bayesian paradigm which enables a complete quantification of uncertainties in date estimates. However, it is sometimes too slow and computationally demanding to be used, especially when large numbers of sequences are involved. Alternatives based on optimisation have therefore started to appear, including LSD (11) which uses least-square optimisation methods and TempEst (12) which uses a linear regression to explore the temporal structure of the data. A systematic comparison between LSD, TempEst and BEAST reported that they produced highly congruent estimates of evolutionary rates (13). More recently, three new optimisation methods have been released based on maximum likelihood, namely node.dating (14), treedater (15) and TreeTime (16). All these methods are faster than BEAST and able to deal with larger datasets, in great part due to the fact that they assume that phylogenetic relationships have previously been assessed. Their input data therefore consists of the sampling dates plus an unrooted phylogenetic tree which needs to be built in a separate analytical step using a standard phylogenetic software such as RAxML (17), PhyML (18), FastTree (19) or IQ-TREE (20).

Here we present a new methodology called BactDating for analysing dated genetic data in order to estimate evolutionary rates and dated phylogenies in bacterial populations. We use a Bayesian framework for inference as in BEAST, but consider that phylogenetic relationships have been assessed in a previous step as in the optimisation and maximum likelihood methods described above. This way we enjoy the benefits of Bayesian inference in ancestor dating (21), such as assessment of uncertainties and flexibility of model choice and comparison, but with a computational scalability and speed comparable to the optimisation methods described above. Furthermore, we explore the specific problems posed by application in bacterial genomics, and in particular the disruptive effect that homologous recombination can have on estimates of the temporal signal (22, 23). Recombination is well known for disrupting phylogenetic inference, and especially to affect branch lengths estimates so that trees look star-like with abnormally long terminal branches (24, 25, 22). To account for this, sites detected as recombinant are sometimes removed prior to running BEAST, but this approach is inefficient and can even exacerbate the problem (22). Instead we show how the effect of recombination can be accounted for in the dating of ancestral nodes, by exploiting a phylogenetic method that accounts for bacterial recombination such as ClonalFrameML (26) or Gubbins (27).

We applied BactDating to a large number of datasets simulated under various conditions in order to benchmark its ability to produce correct estimates by comparison with the correct parameter values used during simulation. We also demonstrate the usefulness of BactDating on three case studies based on real datasets from recently published bacterial genomic studies. The first case study used ancient DNA sequencing in order to compare medieval and modern genomes of the leprosy causing pathogen *Mycobacterium leprae* (28). In the second case study a large number of isolates from clonal lineage of *Shigella sonnei* from Vietnam were sequenced and compared to study local emergence and dissemination (29). Finally, in the third case study, a worldwide collection of genomes from a highly recombining lineage of *Streptococcus pneumoniae* were used to investigate its global success and spread (30).

## MATERIALS AND METHODS

### Overview of Bayesian inference

We consider as input a phylogenetic tree 𝒫 previously estimated from a set of *n* bacterial genomes using a standard phylogenetic method. For ease of presentation, we initially make two simplifying assumptions that will be relaxed later. Firstly, we consider that all the isolation dates of the genomes are known. Secondly, we assume that the tree 𝒫 is already rooted, so that it contains *b* = 2*n* − 2 branches. Our aim is to estimate a dated tree 𝒯, which in this case means estimating the dates at which each of the *n* − 1 internal nodes in 𝒫 existed. There are two key differences between the input phylogeny 𝒫 and the target of inference, the dated or time-calibrated tree 𝒯. First, the branch lengths of 𝒫 are measured in units of the expected number of substitutions, whereas the branch lengths of 𝒯 are measured in calendar time. Second, as a consequence, the ‘heights’ of all tips and internal nodes in 𝒯 are directly interpretable as calendar dates, which is not true of 𝒫.

To estimate the dated tree 𝒯 in a Bayesian inferential framework, we need to specify a prior on 𝒯 and the likelihood of observing the substitutions in 𝒫 given the dated tree 𝒯. For this likelihood, we will consider three models of increasing complexity: a strict clock model without recombination, a relaxed clock model without recombination and finally a strict or relaxed clock model with recombination. The main notation is summarised in Table 1.

**Table 1:** Table of symbols

| Symbol | Description |
| --- | --- |
| $\mathcal{P}$ | Input phylogenetic tree |
| $n$ | Number of leaves in the phylogenetic tree $\mathcal{P}$ |
| $b$ | Number of branches in the phylogenetic tree $\mathcal{P}$ |
| $x_i$ | Length of the $i^{\text{th}}$ branch of the phylogenetic tree $\mathcal{P}$ |
| $\mathcal{T}$ | Dated phylogeny to be estimated |
| $l_i$ | Duration of the $i^{\text{th}}$ branch of the dated phylogeny $\mathcal{T}$ |
| $\Theta$ | Additional parameters to be estimated |
| $\alpha$ | Coalescent time unit |
| $\mu$ | Mean substitution rate |
| $m_i$ | Substitution rate for the $i^{\text{th}}$ branch of the dated phylogeny $\mathcal{T}$ |
| $\sigma$ | Standard deviation of the per-branch substitution rates |

More formally, we want to jointly infer the dated genealogy 𝒯 and some additional model parameters Θ given an estimated phylogeny 𝒫, so that the target distribution is:
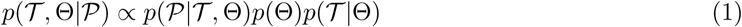

The first term 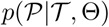 is the likelihood, which is described in subsequent sections under various conditions. The second term *p*(Θ) represents the prior on the additional parameters in Θ and will also be described later. The third term 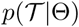 is the prior on the dated genealogy 𝒯 for which we consider a coalescent model with constant population size (31), which is the genealogical process that corresponds to many forward in time population genetics model such as the standard neutral Wright-Fisher model. The only parameter of this model is the coalescent time unit *α* = *N_e_g* which is the product of the effective population size *N_e_* and generation time *g*. The parameter *α* is included in the vector Θ of parameters that we aim to co-estimate. This prior term 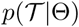 can be computed by considering the ordered list of 2*n* − 1 times *t_i_* of both terminal and internal nodes in the dated genealogy, and the values *k_i_* of lineages existing in each time interval, which gives (32):
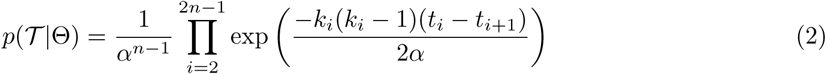

### Strict clock model

We break down the likelihood 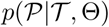 into the product of the individual likelihoods of the observed number of substitutions, *x_i_*, on each branch *i* ∊ {1,…, *b*} of the input phylogeny 𝒫 given the duration, *l_i_* of that branch in the dated tree 𝒯. Substitution models typically consider a discrete number of substitutions on each branch. For example in the strict clock model (33) the same rate *µ* of evolution is applied to all branches, so that the number of substitutions *x_i_* is simply distributed as *x_i_* ~ Poisson(*µl_i_*), where *x_i_* is discrete. However, phylogenetic software typically estimate the branch lengths *x_i_* as a continuous variable, due in particular to the use of non-homogenous mutation models (34) and uncertainties in phylogenetic reconstruction (35). Consequently, we consider here a Gamma distribution, with mean equal to its variance by analogy with the Poisson distribution, so that the likelihood function becomes:
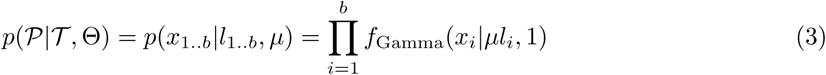

where the rate *µ* is included in the vector of parameters Θ. The Gamma distribution used above and throughout this article is parameterized in terms of the shape and scale parameters, respectively.

### Relaxed clock model

In practice the assumption of a strict clock rate may be inappropriate, so next we consider an uncorrelated relaxed clock model where each branch has a specific rate *m_i_* sampled from a given distribution (36). For example this distribution could be *m_i_* ~ Gamma(*k, θ*), so that the product of the rate *m_i_* and the branch length *l_i_* is distributed as *m_i_l_i_* ~ Gamma(*k, l_i_θ*). If we now consider substitution as a Poisson process with rate *mi_l_i* we find that the number of mutations *x_i_* is discrete and distributed as 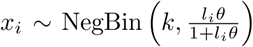 which is the relaxed clock model used by treedater (15).

More generally, let us consider that the per-branch rates *m_i_* are independent and identically distributed samples from an unspecified distribution with expectation and variance respectively equal to **E**(*m_i_*) = *µ* and **V**(*m_i_*) = *σ*^2^. We also allow continuous values for *x_i_* and consider, as we did for the strict clock model in Equation 3, that *x_i_* ~ Gamma(*m_i_l_i_,* 1). By application of the laws of total expectation and variance, we can then deduce the expectation and variance of *x_i_*:
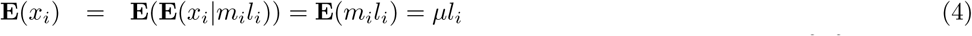

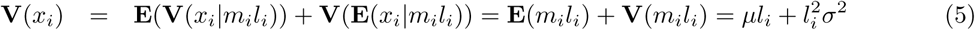

By analogy with the case of the strict clock model in Equation 3, we impose a Gamma distribution with this mean and variance, resulting in the following likelihood function:
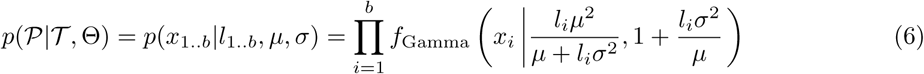

where both *µ* and *σ* are included in the vector of parameters Θ. We note that the special case where the variance of the branch-specific rates is zero corresponds to the strict clock model, so that setting *σ* = 0 in Equation 6 gives Equation 3. This relaxed clock model is similar to the uncorrelated lognormal relaxed clock model (36) implemented in BEAST (3), in the sense that both the mean and the variance of the per-branch rates are independent parameters, whereas a model similar to the uncorrelated exponential relaxed clock model (36) could be obtained by setting *µ* = *σ*^2^. Note however that unlike these previous relaxed models we did not specify a distribution for the per-branch rates, but instead we specified a Gamma distribution for the resulting branch lengths in Equation 6.

### Accounting for bacterial recombination

The input phylogeny to be dated may be the output from phylogenetic software that accounts for the effect of bacterial recombination, for example ClonalFrameML (26) or Gubbins (27). In this case, the output contains for each branch *i* the proportion *c_i_* of the genome that has been found to be non-recombinant on that branch, as well as the recombination-corrected length *x_i_* of each branch. The branch length estimate in 𝒫 is related to *s_i_*, the number of substitutions observed in the *non-recombinant* portions of the genome, and *c_i_* by the formula *x_i_* = *s_i_*/*c_i_*. Such a recombination-corrected phylogeny could be dated as if it were the output of standard phylogenetic software but that may underrepresent uncertainty in the dating because only partial sequence was used to estimate *x_i_*, especially when the fractions 1 − *c_i_* of recombinant material are large. Instead, we implemented dating of such trees based on a modified likelihood function that accounts for the fact that only the non-recombinant regions are informative about the branch lengths. This is achieved by considering the distribution of the number *s_i_* = *x_i_c_i_* of substitutions in the non-recombinant regions and scaling down the substitution rates by a factor *c_i_*. For example, in the case of a relaxed clock model, both *µ* and *σ* are scaled down by a factor *c_i_* so that the likelihood in Equation 6 is modified to give:
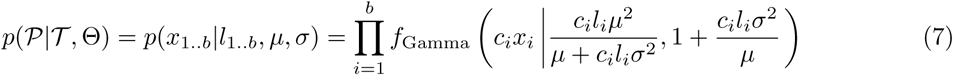

As before, the case of a strict clock is obtained by setting *σ* = 0 in Equation 7, so that the shape and scale parameters of the Gamma distribution become simply *c_i_l_i_µ* and 1, respectively. We have implemented functions that can read directly the output files of ClonalFrameML (26) and Gubbins (27) in order to date recombination-corrected phylogenies using this approach.

### Markov Chain Monte Carlo methodology

We sample from the posterior distribution in Equation 1 using a Markov Chain Monte Carlo (MCMC). Most parameters, such as the age of each node in the dated genealogy 𝒯 are updated using Metropolis-Hastings moves with normal proposals centred on the current value. One exception is the coalescent time unit *α* for which a Gibbs move is available, by noticing that in Equation 2 the rate 1*/α* admits a Gamma conjugate prior. Specifically, we consider a Gamma(*k, θ*) prior on 1*/α*, so that the posterior distribution of *α* is distributed as:
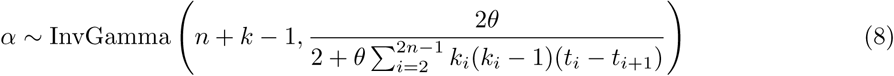

The priors on the parameters *µ*, *σ* and 1/*α* are Gamma(0.001,1000) by default and in all applications below.

We have so far been assuming that the root of the phylogeny 𝒫 was predetermined for example using one or several outgroup sequences, and also that all sampling dates of the genomes in 𝒫 were known exactly. However, both of these assumptions can easily be relaxed via data augmentation in which the location of the root in 𝒫 and the unknown sampling dates are treated as additional parameters co-estimated using additional MCMC moves (37). For the location of the root, we consider as a prior that all points on the phylogeny are equally like to be the root and we use two Metropolis-Hastings moves, one proposing to move the root from its current location to one of the branches directly underneath, and another proposing to move the root while staying on the same branch. For the sampling dates, the user can specify the bounds of the uniform prior considered as possible dates, or by default the range of all known sampling dates is used, and a Metropolis-Hastings move proposes to update the unknown sampling dates within their allowed range.

Options are available to perform inference under the strict clock model (Equation 3) or under the relaxed clock model (Equation 6), but by default we consider a mixture of the two models, in which half of the prior weight is given to each model. Mixing between the two models is implemented using reversible jumps to propose moves between the strict (*σ* = 0) and relaxed (*σ* > 0) models (38). This allows us to perform model comparison between the two models, and in particular to estimate the Bayes Factor as the ratio of MCMC iterations spent in each model (39). In summary, each MCMC iteration consists of the following MCMC moves, all of which are used by default but can be deactivated by the user:

- A Metropolis-Hastings move proposing to update the value of the mean substitution rate *µ*
- A Gibbs move updating the coalescent unit *α*
- When using the relaxed clock model, a Metropolis-Hastings move proposing to update the standard deviation *σ* of the per-branch substitution rates
- A reversible-jump move proposing to move from the strict clock model to the relaxed clock model or vice-versa
- For each internal node of the tree, a Metropolis-Hastings move proposing to update its date
- For each leaf of the tree with unknown sampling date, a Metropolis-Hastings move proposing to update its date
- Two Metropolis-Hastings moves proposing to update the root location

By default, the MCMC is run for a total of 10^5^ iterations, with the first half discarded as MCMC burnin and the remainder sampled every 100 iterations. For all results presented below, the convergence and mixing of the chains was assessed using the R package coda (40). The effective sample size of the inferred parameters *α*, *µ* and *σ* were computed to make sure that they were greater than 200. Furthermore, multiple chains were run separately and compared to ensure that the multivariate version of the Gelman-Rubin diagnostic (41, 42) was lower than 1.1.

### Implementation

The methodology described above was implemented in a new R package called BactDating and freely available at https://github.com/xavierdidelot/BactDating. For maximum computational efficiency, the likelihood and prior functions described in Equations 2-7 were written in C++ and integrated into the R package using Rcpp (43). BactDating also includes functions to simulate dated coalescent trees from Equation 2, and phylogenetic trees from Equations 3 and 6, which we used to simulate datasets and assess the performance of our inference methodology.

BactDating also includes a function to perform root-to-tip linear regression analysis, including optimisation of the root to maximise the coefficient of determination *R*^2^, and implementation of a previously described test to assess the significance of the temporal signal based on random permutations of sampling dates (44). This linear regression procedure is used to provide a good default starting point for the MCMC algorithm. Finally, several studies have proposed that the significance of the temporal signal can be tested by comparison with a run where all sampling dates are set equal (45, 1, 46, 47), and we implemented this approach by computing the deviance information criterion DIC (48) for the two runs with and without sampling dates set equal.

## RESULTS

### Application to a single simulated dataset

To demonstrate the use of our Bayesian methodology, we first simulated a single dataset, consisting of 100 individuals, sampled at regular intervals between the year 2000 and 2010. The genealogy was drawn from the heterochronous coalescent model (Equation 2) with coalescent time unit equal to *α* = *N_e_g* = 5 years (Figure 1A). The strict molecular clock model (Equation 3) was applied to this genealogy with mean rate of *µ* = 5 substitutions per year to obtain an unrooted phylogenetic input tree (Figure 1B). We also consider the sampling dates as part of the input, except that each individual had a 10% probability of having an unknown sampling date. We first performed a linear regression analysis of root-to-tip distance versus sampling dates (when known), with the root position selected to optimise temporal signal. This resulted in a slightly underestimated clock rate of *µ* = 4.38 substitutions per year, and a root located on the correct branch as in Figure 1A, but with an estimated date of 21 February 1996, underestimated compared to the correct root date 28 December 1996. This linear regression had a high fraction of variance explained by the model, *R*^2^ = 0.86, with all points falling within or very close to the 95% confidence intervals (Figure 1C), and a highly significant p-value of *p* < 10^−4^ based on a permutation test (44).

**Figure 1:**
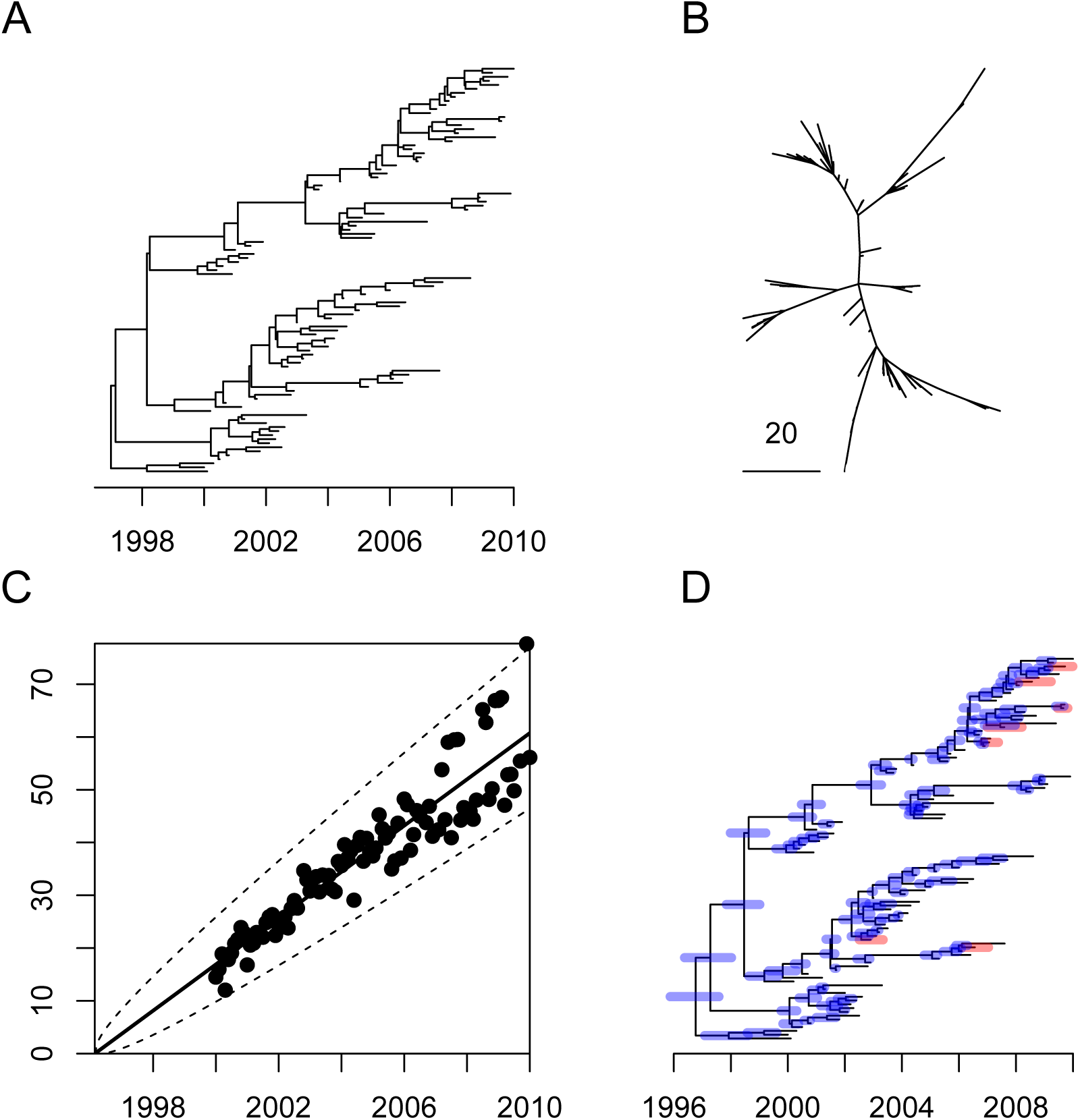
Application to a single simulated dataset. (A) the correct dated genealogy. (B) the unrooted phylogeny used as input. (C) Linear regression of root-to-tip (y-axis) versus sampling dates (x-axis). (D) Estimated dated genealogy, with blue bars indicating 95%CI for ancestral dates and red bars representing the 95%CI for the unknown sampling dates.

The clock rate and tree root estimated by the linear regression were both used as starting point for our MCMC procedure. The run time for the default 10^5^ iterations was about 10 minutes on a standard desktop computer. Values in square brackets below represent the 95% credible intervals (95%CI) of estimated parameters. The posterior distribution of the coalescent time unit *α* had mean 4.69 years [3.66-5.98], which includes the correct value of 5 years used in the simulation. The substitution rate *µ* had mean 4.96 per year [4.46-5.47], which also includes the correct value of 5 per year. The posterior probability of the root location was highest for the correct branch, but only equal to 0.56 with the remaining probability being shared between the two branches directly below the short branch stemming from the real root (Figure 1A). Because of the shortness of this branch it is not surprising that there is uncertainty about the exact location of the root. Posterior mean and 95%CI were also estimated for the dates of all ancestral nodes and leaves for which the sampling dates were unknown (Figure 1D). In particular, the root of the tree had a mean date 24 September 1996 [30 October 1995 - 4 August 1997] which covers the correct date 28 December 1996.

### Application to multiple simulated datasets

We repeated the procedure described above for 100 simulated datasets, each of which was generated with the same coalescent time unit *α* = 5 years but with the substitution rate *µ* varying between 0.1 and 10 per year. For each dataset, we estimated the mean and 95%CI of the two parameters *α* and *µ* (Figure 2A). We found that estimated values for *α* remained around the correct value of 5, with most 95%CI covering 5, whereas the estimates of *µ* increased with the correct value of *µ*, with once again most 95%CI covering the correct values. We then repeated the procedure again for another 100 simulated datasets, but this time keeping *µ* = 5 fixed and varying *α* between 0.1 and 10 year. As expected, we found that in these conditions the estimated values of *µ* remained constant and that the estimated values of *α* followed the correct values used in the simulations (Figure 2B).

**Figure 2:**
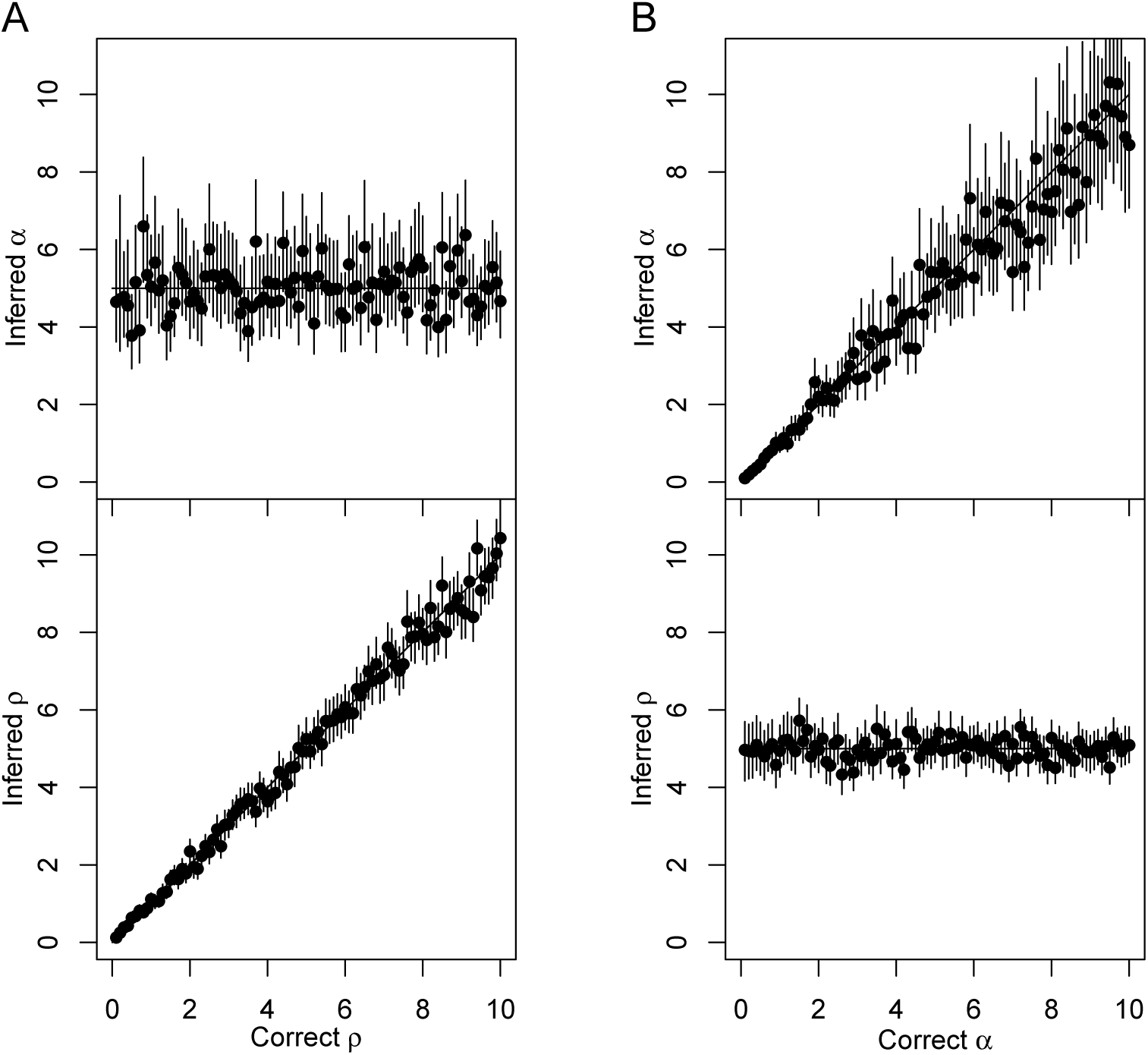
Application to multiple datasets simulated with a strict clock. (A) One hundred simulated datasets were analysed, each of which used parameters *α* = 5 and 0.1 < *µ* < 10 (x-axis), and for both parameters the inferred mean (y-axis, dot) and 95%CI intervals (y-axis, line) are shown. (B) Same as panel A, but using a different set of one hundred simulations for which the true parameters were 0.1 < *α* < 10 (x-axis) and a fixed *µ* = 5.

The simulations considered so far were generated using a strict molecular clock (Equation 3) and inferred using a 50-50 mixture of the strict and relaxed clock models (cf Methods). The inferred Bayes Factors were always overwhelmingly in favour of the correct strict model, with the exception of only the first two simulations in Figure 2A, for which *µ* = 0.1 and *µ* = 0.2 substitutions per year, respectively. The strict clock rate used in these simulation was too low to rule out a relaxed clock model, and doing so would require a sampling interval of more than 10 years. We now consider a new set of 100 simulations performed under the relaxed clock model (Equation 6), in which the coalescent time unit is *α* = 5, the average rate is *µ* = 5 per year, and the standard deviation of the clock rate *σ* varies between 0.1 and 10. Inference was performed once again under the mixed model, in exactly the same conditions are previously. The estimates of the coalescent unit *α* and the average clock rate *µ* remained around the correct value of 5, with most 95%CI covering this value, but we note that as the standard deviation *σ* increased, so did the uncertainty on *µ* (Figure 3). The inferred values of *σ* followed the correct values, except when *σ* was smaller than 2, in which case *σ* was often inferred to be zero (Figure 3). This corresponds to datasets in which the model was incorrectly inferred to be the strict clock model (*σ* = 0) instead of the relaxed clock model (*σ* > 0). This behaviour is expected, since when the standard deviation *σ* of the per-branch clock rates is small (relative to its mean *µ*) the relaxation of the clock has little effect and therefore the data is hard to differentiate from data generated under the strict clock model. This incorrect model selection is therefore not an issue, and other parameter estimates such as the coalescent time unit *α* and evolutionary rate *µ* are unaffected (Figure 3). However, this behaviour demonstrates that our algorithm is relatively conservative in calling the clock relaxed, as a result of our choice of a highly uniformative prior on *σ* in the relaxed clock model which has a direct impact on model selection (49).

**Figure 3:**
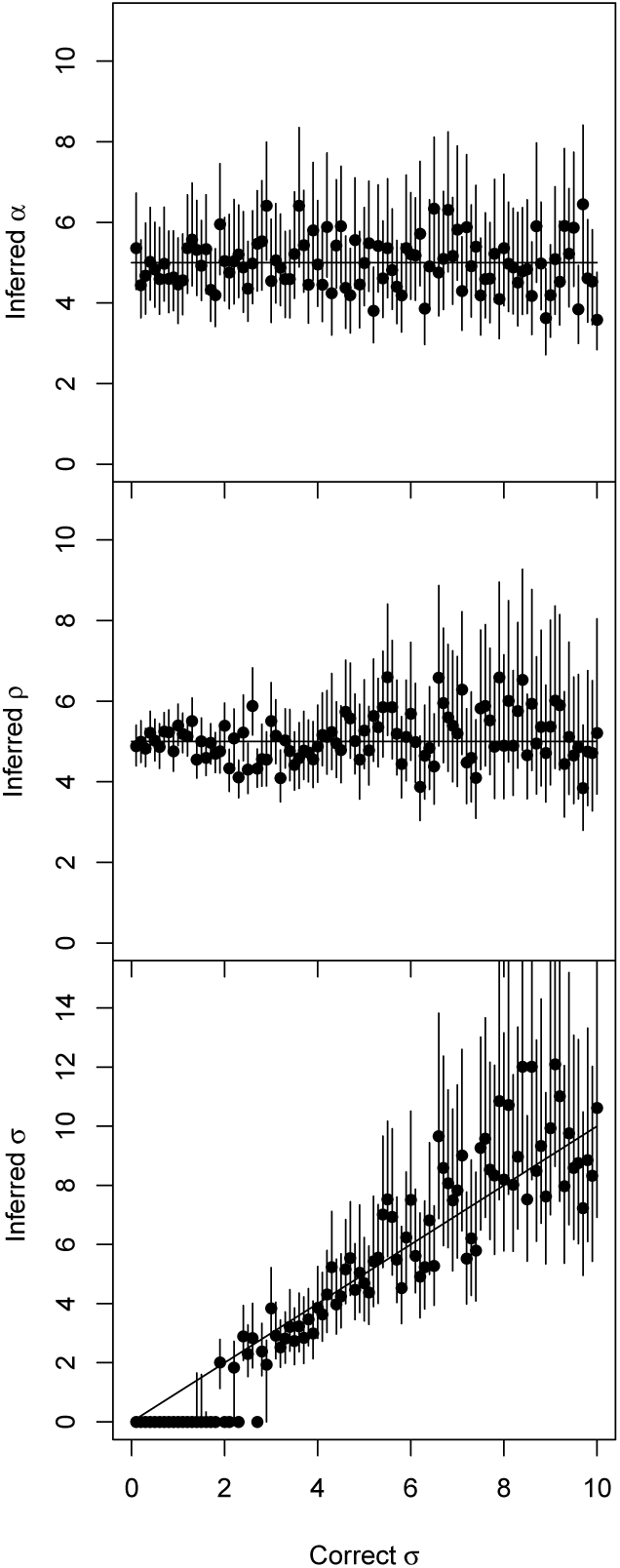
Application to multiple datasets simulated with a relaxed clock. One hundred simulated datasets were analysed, each of which used parameters *α* = 5, *µ* = 5 and 0.1 < *σ* < 10 (x-axis), and for these parameters the inferred mean (y-axis, dot) and 95%CI intervals (y-axis, line) are shown.

Taken altogether, these results on simulated data indicate that our MCMC procedure is correct, and that there is significant statistical power to estimate the key parameters of the models, and therefore to accurately perform Bayesian inference on the ancestral dates of a phylogeny, at least in the conditions used for simulating these datasets. The range of parameters used in the simulations above were selected to be representative of typical situations that arise in the genomic epidemiology of bacterial populations. In particular, the genome-wide substitution rate varies between species in the same order of magnitude considered above between 0.1 and 10 substitutions per year (50, 2, 23). Sequencing a sample of 100 genomes is also frequently achievable nowadays thanks to the recent reduction in cost and time required to sequence whole bacterial genomes (51). The assumption of a uniform unbiased sampling frame over 10 years represents a good case scenario, which is not always achievable. When it is not, the statistical power to accurately date a phylogenetic tree is likely to be reduced, and therefore the uncertainty in reconstructions is increased, which our Bayesian method is well suited to capture.

### Application to an ancient bacterial pathogen using aDNA

*Mycobacterium leprae* is the causative agent of leprosy, a debilitating disease that was endemic throughout Europe in the Middle Ages, and still remains a critical health threat in some parts of the developing world (52). Here we reanalyse previously published data from (28) including ten recent genomes (sampled between 1982 and 2012) and five ancient genomes (sampled between 990 and 1369). An unrooted phylogenetic tree was reconstructed using PhyML (18) (Figure S1). After selecting the root that maximises the coefficient of determination *R*^2^ = 0.9, we find a strong correlation between sampling dates and root-to-tip distances (Figure 4A), with an estimated rate of 0.0353 substitution per genome per year and estimated root date of 928 BCE. All root-to-tip distances fall within the interval expected under a strict molecular clock (Figure 4A) and despite the low number of tree leaves, a date randomisation test (44) found that the temporal signal is significant (*p* < 10^−4^).

**Figure 4:**
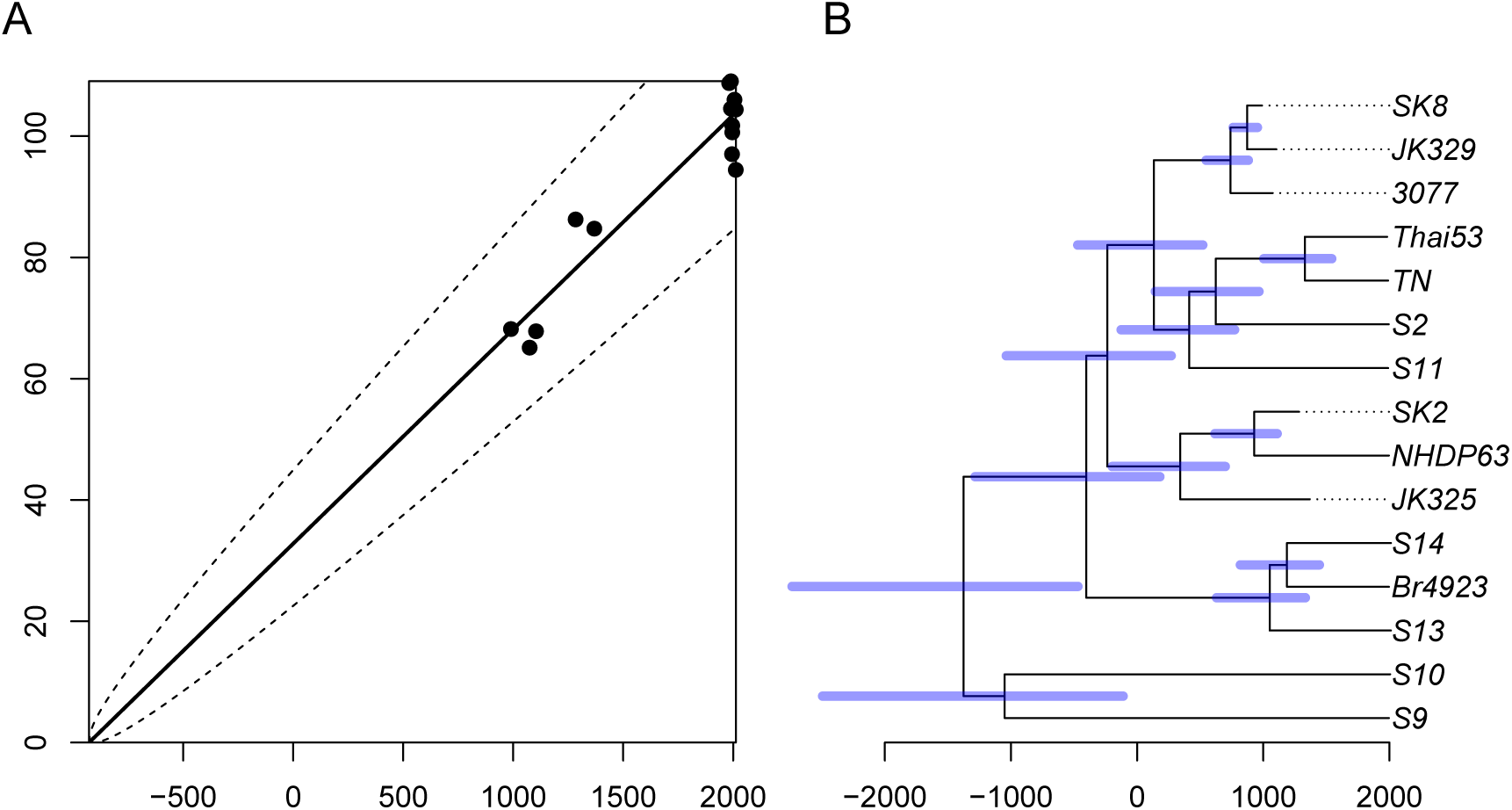
Analysis of *Mycobacterium leprae* dataset. (A) Linear regression of root-to-tip (y-axis) versus sampling dates (x-axis). (B) Estimated dated genealogy, with blue bars indicating 95%CI for ancestral dates.

We performed the default 10^5^ MCMC iterations, which took less than a minute to run. The dated phylogeny produced (Figure 4B) has the same root as for the root-to-tip analysis above, with mean dating of 1396 BCE and a broad 95%CI of [2735-490] BCE (Figure 4B). A strict clock model was inferred, with a Bayes Factor of 141.85. The clock rate had a posterior mean of 0.0314 substitutions per genome per year [0.0219-0.0419] (Figure S2). These estimates are in excellent agreement with the original analysis of this data using BEAST (28). The substitution rate is low compared to values reported in similar bacterial phylogenomic studies as was previously reported (28, 23), which is probably a result of both a low mutation rate in *M. leprae* and the negative dependency between substitution rate estimates and sampling time (53, 2, 23). To test further the significance of the temporal signal in this dataset, the MCMC was rerun under the assumption that all genomes were sampled on the same date. The deviance information criterion DIC (48) in this run was 243.28 compared to 170.57 when the correct dates were used, which indicates conclusively that the temporal signal is significant (45).

### Application to a locally emerging clonal bacterial lineage

The four *Shigella* species are Enterobacteriaceae that have adapted to a human-restricted pathogenic lifestyle and become some of the most prevalent causes of human dysentery (54). The recent spread of antibiotic resistant lineages of *S. sonnei* to several developing countries where *S. sonnei* is traditionally rate is a major global health concern (55, 56). We reanalysed previously published genomic data on the spread of the VN clade in Vietnam (29). *S. sonnei* is a clonal species, with only a single recombination event reported in a species-wide genomic study (55). No recombination event was reported in the VN dataset (29) and a ClonalFrameML (26) analysis found no recombination event either. A phylogenetic tree was constructed using PhyML (18) using 161 whole genomes sampled from Ho Chi Minh City (Vietnam) between 1995 and 2010 (Figure S3). This phylogeny contained six outgroup genomes which were used to establish the location of the root for the remaining genomes (Figure S3). As previously reported (29), the correlation between root-to-tip distances and isolation dates is very strong with a coefficient of determination *R*^2^ = 0.91, and this result was found to be statistically significant according to a randomisation test (*p* < 10^−4^, Figure S4). This linear regression suggests a clock rate of 3.74 and a root date of 1982.68.

Running our algorithm for the default 10^5^ MCMC iterations on this dataset took about ten minutes on a standard computer. Since only the year of the isolation dates were known, we allowed them to vary using a uniform prior within that year. The resulting dated phylogeny (Figure 5A) has mean dating 14 June 1983 [28 December 1977 - 18 November 1986], which is in excellent agreement with the previous report based on BEAST of 1982 [1978-1986] (29). A relaxed clock model was selected with a Bayes Factor greater than 1000 against the strict clock model. The inferred substitution rates had mean *µ* = 4.22 substitutions per year [3.66-4.85]. This is equivalent to 8.34 × 10^−7^ [7.24 × 10^−7^-9.59 × 10^−7^] substitutions per site per year, which is in excellent agreement with the previous estimate from BEAST of 8.5 × 10^−7^ [7.6 × 10^−7^-9.5 × 10^−7^] (29).

**Figure 5:**
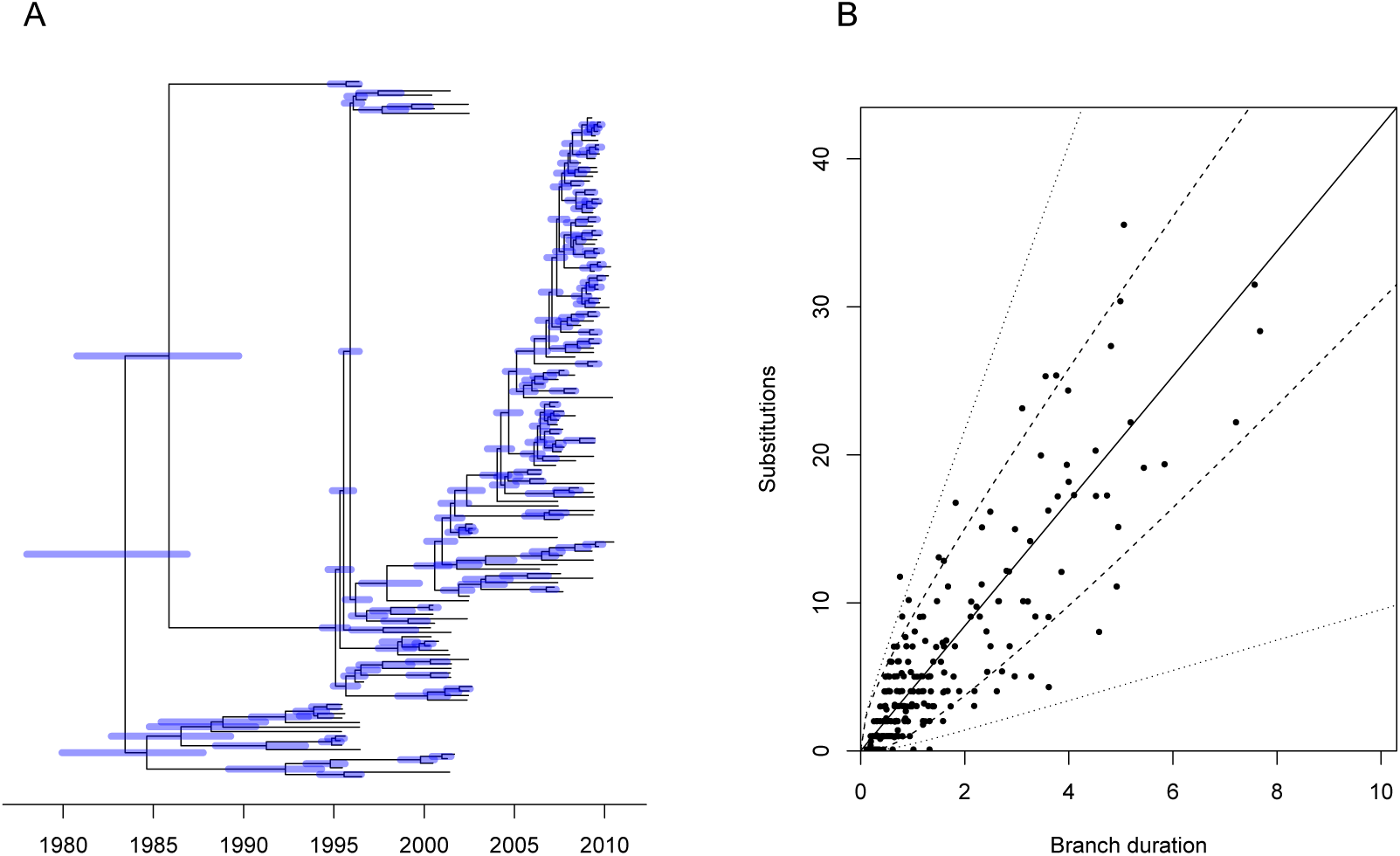
Analysis of *Shigella sonnei* VN dataset. (A) Estimated dated genealogy, with blue bars indicating 95%CI for ancestral dates. (B) Branch-by-branch comparison of duration in years (x-axis) and number of observed substitutions (y-axis). The expectation of the clock model is represented by the solid line, the 95% interval for the strict clock model is represented by the dashed lines and the 95% interval of the relaxed clock model is represented by the dotted lines.

The per-branch standard deviation of the relaxed clock model rate was estimated to be *σ* = 2.24 [1.57-3.09]. This is relatively high especially given that in the root-to-tip analysis almost all the genomes were within the 95% intervals expected under a strict clock model (Figure S4). However, such a root-to-tip analysis is not a statistically powerful way of ensuring the validity of a strict clock model, because the root-to-tip distances are not independent of each other. To illustrate the inadequacy of a strict clock model, the number of substitutions on each branch was considered as a function its duration, along with the 95% ranges expected under both the strict clock and relaxed clock model (Figure 5B). Several branches have numbers of substitutions that fall outside of the strict clock range but within the relaxed clock range, illustrating the better fit of the relaxed clock model compared to the strict clock model.

### Application to a recombining bacterial lineage

*Streptococcus pneumoniae* is a nasopharyngeal commensal and respiratory pathogen of humans, causing a high burden of bacterial pneumonia, sepsis and meningitis worldwide. Originally detected in Spain, the PMEN1 lineage was one of the first multidrug-resistant *S. pneumoniae* found to have spread to multiple continents, and by the late 1990s was responsible for around 40% of infant penicillin-resistant pneumococcal disease in the USA (57). Here we reanalyse previously published genomic data from 238 isolates (30), sampled between 1984 and 2008, although the sampling date was missing for 20 genomes. A phylogenetic tree uncorrected for recombination was constructed using RAxML (17) (Figure S5) and a tree corrected for recombination was built using Gubbins (27) (Figure S6). It was previously reported that correcting for recombination improved the temporal signal, and applying BEAST to the non-recombinant regions resulted in a PMEN1 root date estimate of 1969 [1958-1977] (30). Indeed, we find a coefficient of determination *R*^2^ = 0.22 for a linear regression of root-to-tip distances against isolation dates based on the uncorrected tree (Figure S7), compared with *R*^2^ = 0.59 for the corrected tree (Figure S8). Performing such a linear regression analysis on the uncorrected tree suggests a clock rate of 9.98 substitutions per year and a root date of 1981, whereas on the corrected tree the clock rate is estimated to be 3.21 substitutions per year and the root date 1971.

To illustrate the importance of accounting for recombination when dating lineages, we applied our MCMC algorithm to both the corrected and the uncorrected trees in exactly the same conditions. Each run took approximately 10 minutes using the default settings. Based on the uncorrected tree, a relaxed clock model was inferred with a mean rate *µ* of 3.72 [2.60-4.91] substitutions per year, and per branch standard deviation *σ* of 5.68 [3.91-7.66]. The higher value of *σ* compared to *µ* indicates that the clock is very relaxed, so that estimated dates are highly uncertain (Figure 6A). The root date for example is estimated to be 1523 with a 95% credible interval covering more than six centuries, from 1219 to 1885. The deviance information criterion DIC (48) was 3226.98 which is comparable to 3286.34 when all sampling dates were assumed identical, which suggests that the temporal signal is not strongly statistically significant in this uncorrected tree (45), even though a permutation test on the root-to-tip analysis (44) suggests it is (*p* < 10^−4^).

**Figure 6:**
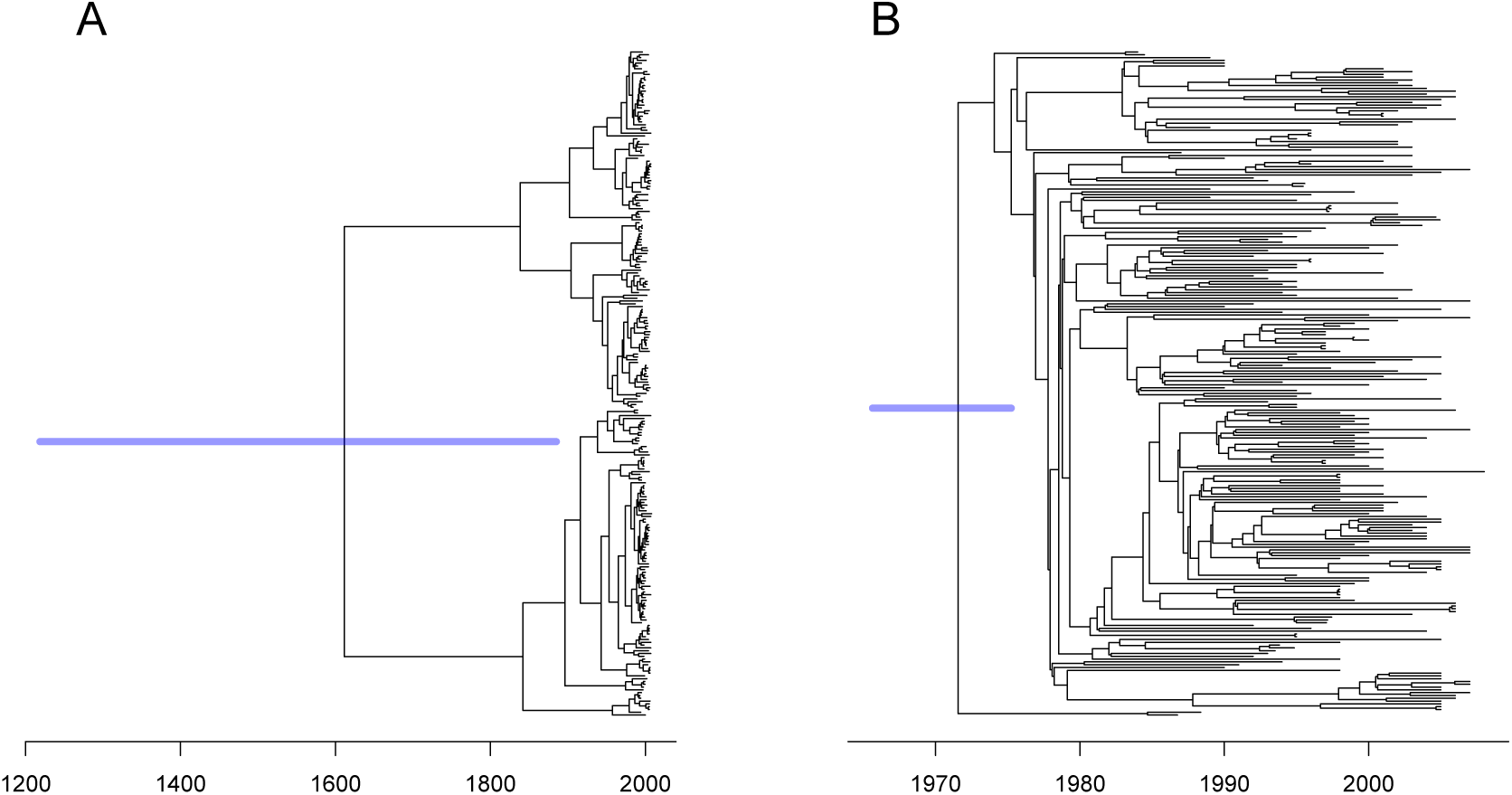
Dating of *Streptococcus pneumoniae* PMEN1 before and after correcting for recombination. (A) Application of dating based on the RAxML tree uncorrected for recombination. (B) Application of dating based on the Gubbins tree corrected for recombination.

When dating was applied to the recombination corrected tree, a relaxed clock model was also selected but this time the mean rate *µ* was 3.09 substitutions per genome per year [2.68-3.53] and the standard deviation *σ* was 1.04 [0.77-1.40]. Thus the clock is less relaxed than for the uncorrected tree, and the dates are more accurate (Figure 6B), for example the date of the root was estimated to be 1972 [1966-1977] which is in excellent agreement with the previous estimate of 1970 based on both root-to-tip analysis and BEAST (30). The deviance information criterion DIC was 3631.94 compared to 6725.77 when all sampling dates were set equal, which suggests that the temporal signal is definitely significant in the recombination corrected tree.

## DISCUSSION

We have presented a new Bayesian approach called BactDating to produce dated phylogenies from a set of bacterial genomes. A key aim was to make sure that our method was fast and scalable to the large numbers of bacterial genomes that can be sequenced thanks to recent improvements in sequencing technologies (51). Several other fast scalable methods have been recently developed (11, 15, 16) but unlike these tools BactDating is based on the Bayesian statistical framework. Bayesian dating provides many advantages (21), such as the ability to naturally quantify uncertainties in parameter estimates, to consider different evolutionary models and to compare them. BactDating is slower than some of these non-Bayesian approaches, but remains fast enough to be applied to datasets of hundreds of genomes in a matter of minutes.

BactDating shares many similitudes with BEAST (3, 4, 5), including the use of a Markov Chain Monte Carlo to perform Bayesian inference, and the applications we presented on three real datasets showed that BactDating and BEAST produce highly consistent results. BactDating is several orders of magnitude faster and more scalable than BEAST, and this is achieved by assuming that the phylogenetic relationships between the genomes have been previously reconstructed using standard phylogenetic software. A first drawback of this approach lies in the computational cost of having to perform this previous analytical step, however this is not a significant issue in practice thanks to the recent development of fast maximum-likelihood phylogenetic software (18, 19, 17, 20) which in most studies are already applied in parallel to dating. A more fundamental drawback concerns the fact that uncertainties associated with phylogenetic reconstruction are not accounted for in the dating. This could be addressed by running BactDating on multiple phylogenetic trees as was proposed in other applications where accounting for phylogenetic uncertainty was a concern (58, 59, 60). The high overall computation cost of this strategy could be avoided through the use of parallel computing, with each node computing for example a bootstrap replicate of the phylogenetic tree and performing dating using BactDating. BEAST explores the full space of unconstrained dated phylogenies, but it should be noted that this creates other issues such as difficulty in MCMC convergence and mixing (61, 62), particularly in the presence of recombination (63), the need to build a consensus tree (64) and the occasional occurrence of non-sensical branches of negative lengths in such trees (65). On the other hand, the use by BactDating of previously assessed phylogenetic relationships can be a significant advantage if the phylogenetic software accounted for the disruptive effect of bacterial recombination, as do ClonalFrameML (26) and Gubbins (27).

Dating phylogenetic events without a prior idea of clock rate is only possible if the temporal signal in the dataset is significant and strong enough (12). This signal is typically assessed using a linear regression of root-to-tip distances versus isolation dates, but this is well known to be problematic since the root-to-tip distances are not independent of one another. Instead, we implemented a previously proposed approach which consists of comparing the results of dating with correct sampling dates and with all sampling dates set equal to one another (45, 1, 46, 47). However, BactDating is also well suited to exploring other options, for example the idea of comparing the results of dating using the correct sampling dates to multiple runs where the sampling dates are randomised (66, 67, 13, 23). So far, this approach has been used rarely in practice because it requires the analysis to be run many times, but our computationally efficient Bayesian framework makes this approach much more applicable than before.

Different substitution models can be used within BactDating as we illustrated by comparing strict and relaxed molecular clock models (Equations 3 and 6) on both simulated and real data. Another extension of the substitution model would be to account for the time dependency of substitution rates. The fact that observed substitution rates are lower on longer time scales compared to recent time scales has been well documented in viral phylogenetics (68, 69, 53) and more recently also in bacteria (2, 23). A model for this dependency, for example an exponential decay equation (53, 23), could be integrated into the distribution of number of substitutions for a given branch in order to test the validity of such a model and to account for this dependency in the dating. A different type of extension would be to consider alternative prior models for the dated phylogeny. Here we assumed a coalescent model with constant population size (Equation 2), but alternatives could easily be implemented such as a skyline model (6, 70). Because it is both Bayesian and computationally efficient, BactDating is well suited to explore and compare such models extensions in future work.

## AVAILABILITY

BactDating is freely available for download from https://github.com/xavierdidelot/BactDating.

